# Genome wide association analysis uncovers variants for reproductive variation across dog breeds and links to domestication

**DOI:** 10.1101/285791

**Authors:** Samuel P. Smith, Julie B. Phillips, Maddison L. Johnson, Patrick Abbot, John A. Capra, Antonis Rokas

## Abstract

The diversity of eutherian reproductive strategies has led to variation in many traits, such as number of offspring, age of reproductive maturity, and gestation length. While reproductive trait variation has been extensively investigated and is well established in mammals, the genetic loci contributing to this variation remain largely unknown. The domestic dog, *Canis lupus familiaris* is a powerful model for studies of the genetics of inherited disease due to its unique history of domestication. To gain insight into the genetic basis of reproductive traits across domestic dog breeds, we collected phenotypic data for four traits, cesarean section rate, litter size, stillbirth rate, and gestation length, from primary literature and breeders’ handbooks. By matching our phenotypic data to genomic data from the Cornell Veterinary Biobank, we performed genome wide association analyses for these four reproductive traits, using body mass and kinship among breeds as co-variates. We identified 13 genome-wide significant associations between these traits and genetic loci, including variants near *CACNA2D3* with gestation length, *MSRB3* and *KRT71* with litter size, *SM0C2* with cesarean section rate, and *HTR2C* with stillbirth rate. Some of these loci, such as *CACNA2D3* and *MSRB3*, have been previously implicated in human reproductive pathologies, whereas others have been previously associated with domestication-related traits, including brachycephaly (*SM0C2*), coat curl (*KRT71*), and tameness (*HTR2C*). These results raise the hypothesis that the artificial selection that gave rise to dog breeds also shaped the observed variation in their reproductive traits. Overall, our work establishes the domestic dog as a system for studying the genetics of reproductive biology and disease.

**Lay Summary:** Variation in reproductive traits across mammals has been extensively investigated and is well established, but the genetic contributors to this variation remain largely unknown. Here, we take advantage of the domestic dog, a powerful model for mammalian genetics, to gain insight into the genetic basis of reproductive traits. By examining the association between more than a hundred thousand genetic variants and four reproductive traits that vary extensively across dog breeds, we identified more than a dozen significant associations for cesarean section rate, litter size, stillbirth rate, and gestation length. Some of the variants that we identify are nearby genes previously implicated in human reproductive pathologies, whereas several others have been previously associated with domestication-related traits. Our results establish the domestic dog as a tractable system for studying the genetics of reproductive traits and underscore the potential for cryptic interactions between reproductive and other traits favored over the course of adaptation.

## Introduction

Mammals exhibit wide variation in traits associated with reproduction (Derrickson 1992, Harrison 2001, Behringer et al. 2006). For example, gestation length can range from 12 days in the Gray dwarf hamster, *Cricetulus migratorius*, to 21 months in the African bush elephant, *Loxodonta africana* (Jones et al. 2009, Kiltie 1992, Martin et al. 1985); neonate size can range from less than one gram in the shrew family (Soricidae), to more than a metric ton in the baleen whales (Balaenopteridae) (Jones et al. 2009, Martin et al. 1985); and neonates can be either precocial (e.g., cricetid rodents, rabbits, and canids) or altricial (e.g., hystricomorph rodents, ungulates, and cetaceans) (Derrickson 1992). This variation in reproductive traits also extends to methods of implantation (Cross et al. 1994), structure of the placenta (Enders et al. 2004, Elliot et al. 2009), and lactation strategies (Pond 1977, Lefèvre et al. 2011). Not surprisingly, many reproductive traits also exhibit substantial intra-specific variation (Kiltie 1982). For example, many mammals exhibit intraspecific variation in gestation length, including primates (Cross et al. 1981), rat and rabbits (Hudson et al. 1999), as well as the domesticated cattle (Burris et al. 1952) and thoroughbred horses (Davies et al. 2002). Similarly, body fat percentages, which are associated with the energetics of reproduction, vary greatly between wild and captive baboons, and intraspecific variation among captive lemurs can vary from 8 – 41% (Dufour et al. 2002).

The existence of phenotypic variation in reproductive traits is well established, and can inform our understanding of the factors that shape patterns of survival and reproduction in both agricultural (Carneiro et al. 2015, Sironen et al. 2010, Bolormaa et al. 2010, Maltecca et al. 2011) and human populations (Walker et al. 2006). Not surprisingly, most genome wide association (GWAS) studies of reproductive traits focus on economically important traits in domesticated species, such as reproductive seasonality in rabbits (Carneiro et al. 2015), infertility in pigs (Sironen et al. 2010), and dairy traits in cattle (Bolormaa et al. 2010). GWAS studies focused on understanding human reproductive biology and its associated pathologies have also shed light on the genetic basis of reproductive traits, including birth weight (Horikoshi et al. 2016) and gestational duration or length (Plunkett et al. 2011, Zhang et al. 2015, Zhang et al. 2017). For example, maternal variation in six genomic loci *{ADCY5, AGTR2, EBF1, EEFSEC, RAP2C*, and *WNT4)* is associated with gestational duration and preterm birth (Zhang et al. 2017). While these studies contribute to our understanding of the genetic architecture of reproductive traits, we still understand very little about the molecular pathways underlying this variation and are unable to explain the majority of the heritability in reproductive traits (Cassady et al. 2001, Langlois et al. 2012, Johanson et al. 2011).

To address this challenge, we studied the genetics of reproductive traits in a powerful new model system: the domestic dog. The dog is well-suited to this question, because the domestication bottleneck followed by intense artificial selection and inbreeding imposed over the past 300 years has led to the generation of more than 340 recognized breeds that exhibit dramatic morphological variation (Neff et al. 2006, Beale and Ostrander 2012, Marsden et al. 2016). Domestic dog breeds also show substantial variation in their reproductive traits. For example, Pomeranians and Norfolk Terriers typically have only 2 pups per litter, whereas Dalmatians and Rhodesian Ridgebacks typically sire 8-9 pups per litter (Evans et al. 2010). Similarly, 80 – 90% of French Bulldogs and Boston Terriers are born via cesarean section due to cephalopelvic disproportion, whereas only 2 – 3% of Australian Shepherds and Shar Peis require cesareans (Evans and Adams 2010). Recent analyses have begun to study the genetic mechanisms that underlie the remarkable morphological variation between modern dog breeds in diverse traits such as snout length, ear erectness, and tail curliness (Hayward et al. 2016, Boyko et al. 2010, Vaysse et al. 2011, Marchant et al. 2017), as well as genetic disease (Karlsson et al. 2008).

To gain insight into the genetic basis of reproductive traits across domestic dog breeds, we collected phenotypic data for four reproductive traits, namely cesarean section rate, litter size, stillbirth rate, and gestation length. We synthesized data from the primary literature and breeders’ handbooks to obtain coverage of between 23 (gestation length) and 97 (cesarean section rate) dog breeds, as well as body mass data from 101 dog breeds. By matching our phenotypic data to genome-wide genotypic data from the Cornell Veterinary Biobank, we performed GWAS analyses and identified 13 genetic loci that are significantly associated with these reproductive traits (using log body mass as a co-variate). Several of these variants are in or near genes previously implicated in human reproduction-related pathologies. The majority of the variants that we discovered to be significantly associated with reproductive trait variation are also associated with domestication-related traits. For example, we found that variation in a gene previously identified to be involved in brachycephaly is also significantly associated with rates of cesarean sections and that variation in genes previously linked to coat phenotypes, such as curliness, is also associated with litter size. These results suggest that selection for breed-specific morphological traits during dog domestication may have also directly or indirectly influenced variation in reproductive traits. More broadly, our results establish the domestic dog as a tractable system for studying the genetics of reproductive traits and underscore the potential for cryptic interactions between reproductive and other traits favored over the course of adaptation.

## Results

To identify SNPs that are significantly associated with four reproductive traits in domestic dog breeds, we conducted across-breed GWAS analyses using a multivariate linear mixed model implemented in the program GEMMA (Zhou and Stephens, 2012). Number of individuals and distribution of breed varied with analysis (Supplementary Table 1). After filtering for MAF (MAF < 0.05; 10,804 SNPs were excluded) and linkage disequilibrium (34,240 additional SNPs were excluded), 115,683 SNPs were included in the GWAS analysis for each reproductive trait. To validate our GWAS approach and analytical choices, we first used our collected values for body mass, a trait whose genetic associations have been previously extensively studied in dogs (36,37). As expected, our analysis recovered the major genes associated with dog breed body mass variation, including *IGF1* (*P* = 2.1 x 10^−31^), *SMAD2(P =* 1.2 x 10^−17^) and *IGF2BP2* (*P* = 5.1 x 10^−11^) (Supplementary Figure 1, Supplementary Table 2).

### Genetic loci that significantly associate with cesarean section rate

To examine whether there is variation in cesarean section rate among breeds, we first identified cesarean section rate values for a total of 97 of the 162 dog breeds with genotypic data (Supplementary Table 1). The cesarean section rate values were derived from a British survey across 151 breeds covering 13,141 bitches, which had whelped 22,005 litters over the course of a 10 year period (Marsden et al. 2016). The frequency of cesarean sections was estimated as the percentage of litters reported to be born by cesarean section. Among the 97 breeds with overlapping genetic data, the median cesarean section rate is 17.1%, with a minimum of 0% in Curly Coated Retrievers and Silky Terriers and a maximum of 92.3% in Boston Terriers (Supplementary Figure 3A).

To identify SNPs that are significantly associated with the observed variation in cesarean section rate across domestic dog breeds, we conducted an across-breed GWAS analysis using 115,683 SNPs and cesarean section values across 95 dog breeds (Figure 1A, Supplementary Figure 2A). As outlined in the *permutation analysis* section of the Methods, we additionally performed a breed-specific permutation and present those variants with a permutation p-value > 0.05 as suggestive associations. We identified three significant SNPs (Supplementary Table S3), two of which mapped to genes, namely paralemmin 3 (*PALM3*, uncorrected *P* = 1.4 x 10^−9^, *Perm P* = 0.001) and sparc-related modular calcium-binding protein 2 (*SMOC2*, uncorrected *P* = 2.0 x 10^−7^, *Perm P* = 0.011), and a third that mapped to the intergenic region between the *CD36* glycoprotein and a lincRNA (uncorrected *P* = 9.7 x 10^−8^, *Perm P* = 0.024) (Figure 1A). Our GWAS analysis also recovered keratin 71 *[KRT71*, uncorrected *P* = 2.9 x 10^−7^, *Perm P* = 0.217), but its permutation p-value was above 0.05.

**Figure 1.**
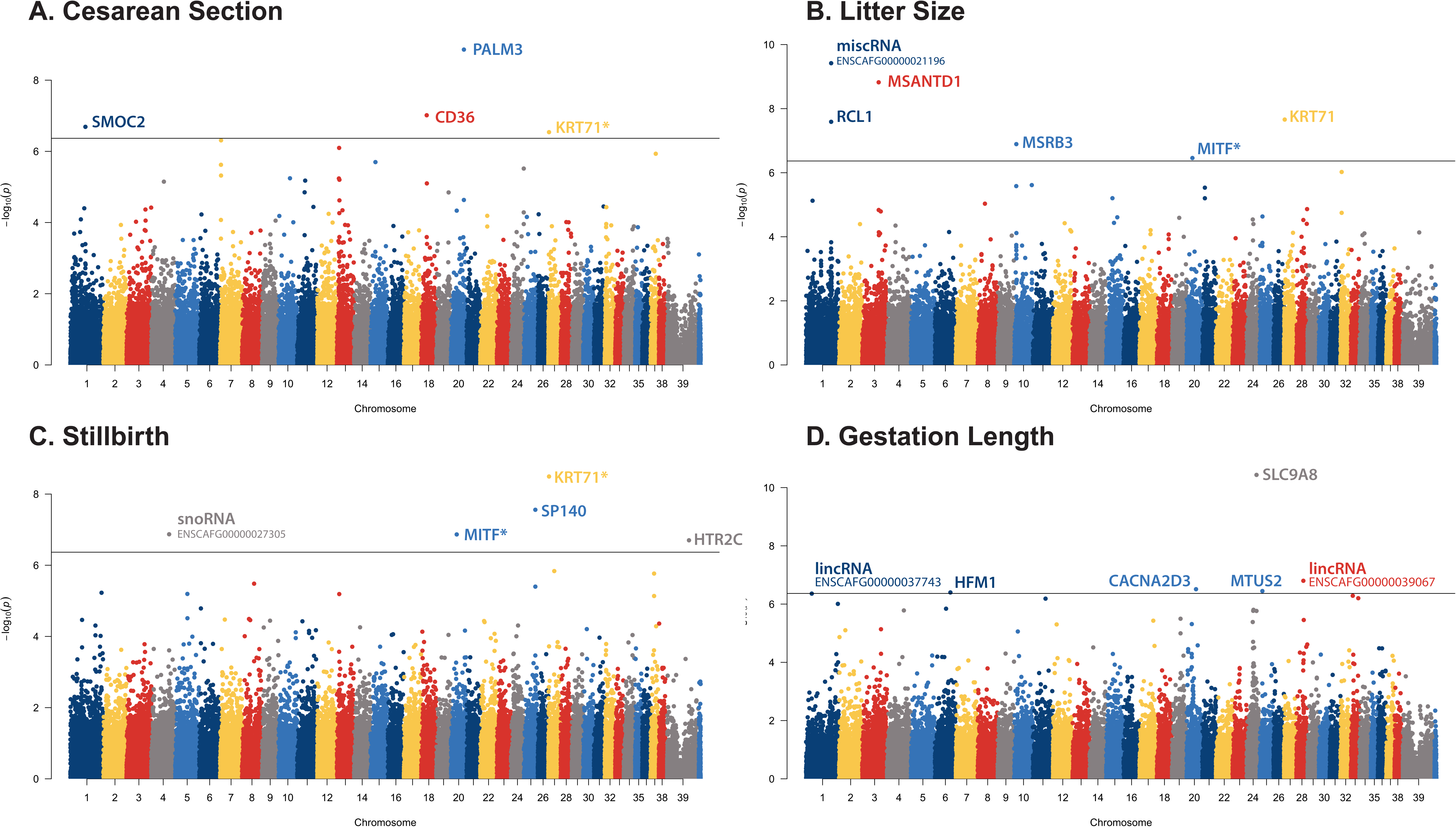
Genome Wide Association results for reproductive traits in domestic dogs. Manhattan plots showing the statistical significance of each SNP as a function of genomic position for (A) cesarean section rate (n = 3,194 individuals, n = 97 breeds), (B) litter size (n = 2,617 individuals, n = 60 breeds), (C) stillbirth (n = 2,590 individuals, n = 57 breeds), and (D) gestation length (n =1,908 individuals, n = 23 breeds). Horizontal line indicates the significance threshold at *P* = 4.3 x 10^−7^. Significant SNPs are labels with the intersecting or nearest gene. Significant SNPs whose permutation p-values were above the 0.05 threshold are indicated by an asterisk (*). Plots were generated in R using the qqman package.

The first significantly associated SNP (chromosome 1: 55,983,871) is found in the intron between exons 13 and 14 of *SMOC2*, a gene that is associated with brachycephaly in dogs (Marchant et al. 2017, Bannasch et al. 2010); variation in *SMOC2* accounts for 36% of facial length variation in dogs (Marchant et al. 2017). In humans, *SMOC2* is highly expressed in endometrium as well as other reproductive tissues, including the fallopian tubes, ovaries and cervix (Figure 2) (Uhlén et al. 2015). The 3’ intronic location of the SNP raises the possibility that it might be regulatory (Rose 2008).

**Figure 2.**
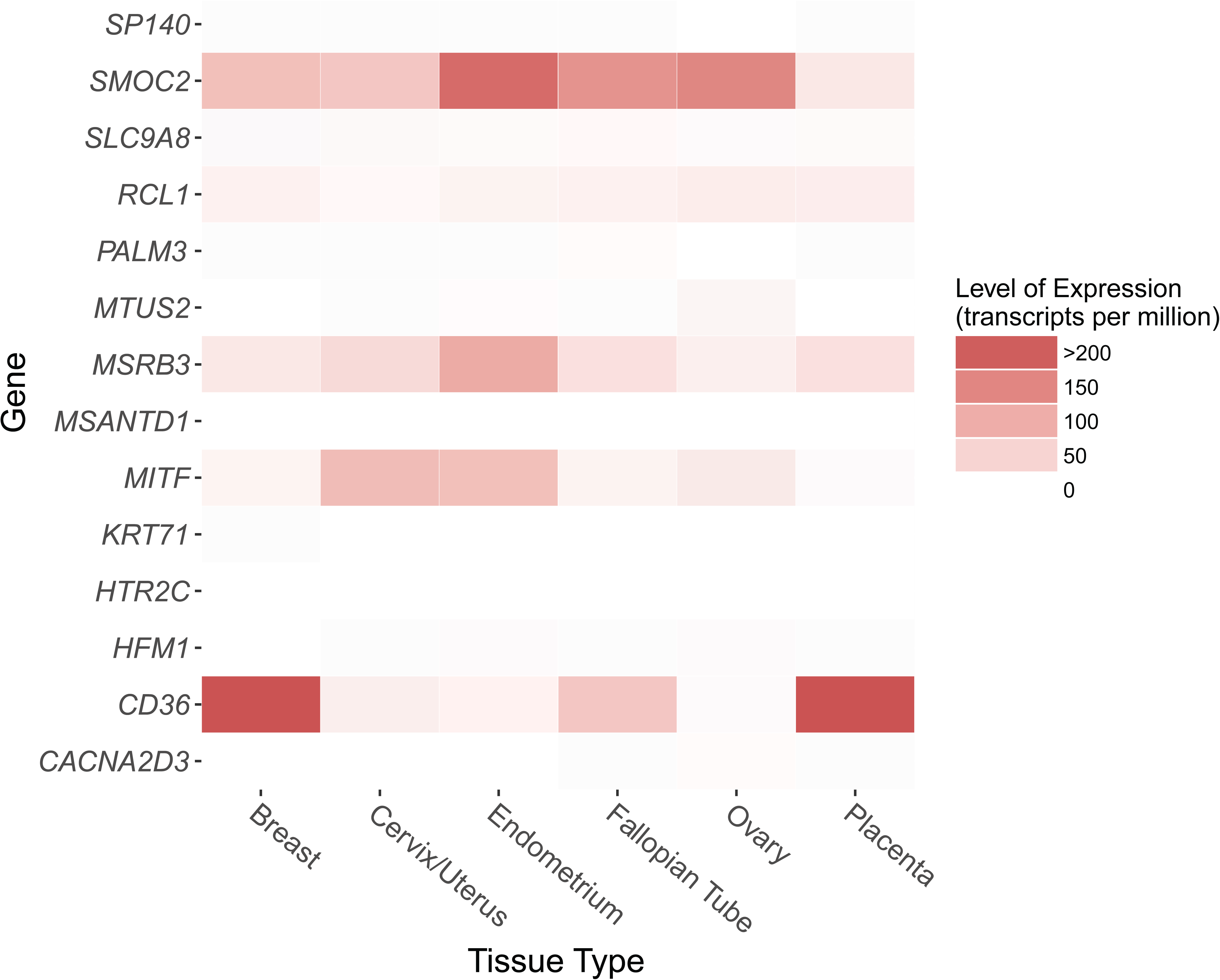
Gene expression in human female reproductive tissues of genes that contain or are adjacent to SNPs associated with reproductive traits in domestic dogs. Raw data were obtained from the Human Protein Atlas database (42).

The second SNP is found in the 3’ UTR of *PALM3*, which is a member of the paralemmin gene family that also includes *PALM1, PALM2*, and *PALMD* (palmdelphin); members of this family are implicated in plasma membrane dynamics and as modulators of cellular cAMP signaling in the brain (Basile et al. 2006, Kutzleb et al. 1998). The function of *PALM3* may be slightly different from the rest of the genes in the family, with recent work suggesting that PALM3 is a binding protein of the single immunoglobulin IL-1 receptor-related molecule (SIGIRR), which is a negative regulator of Toll-Interleukin-1 receptor signaling (Chen et al. 2011). In humans, *PALM3* is primarily expressed in the membranes of the stomach, kidney, parathyroid gland and epididymis (Figure 2) (Uhlén et al. 2015). The SNP (chromosome 20: 48,454,259) that is significantly associated with cesarean section rate is found in the first intron of the *PALM3* gene, suggesting that it might be involved in regulatory actions typically observed in 5’ introns (Rose 2008).

The third significant SNP (chromosome 18: 20,272,961) is found in the intergenic region between the *CD36* gene and alincRNA (ENSCAFG00000034312). The protein product of CD36 is the fourth major glycoprotein of the platelet surface and serves as a receptor for thrombospondin in platelets (Tandon et al. 1989). Other known functions include transport of long chain fatty acids (Febbraio et al. 2001). However, we believe that this variant is more likely a replication of a previous association between the nearby *FGF4* retrotransposon and chondrodysplasia across breeds (Parker et al. 2009).

### Genetic loci that significantly associate with litter size

To examine whether there are SNPs that are significantly associated with variation in litter size among breeds, we retrieved litter size data from 10,810 litters of 224 breeds registered in the Norwegian Kennel Club (Borge et al. 2012). For these data, we were able to obtain average number of pups per litter values for 60 of the 162 dog breeds with overlapping genetic data (Supplementary Table 1). Among these 60 breeds, median litter size is 5.55 pups, with a maximum 8.9 in Rhodesian Ridgebacks and a minimum of 2.4 in Pomeranians (Supplementary Figure 3B).

To identify SNPs, and genes proximal to them, that are significantly associated with the observed variation in litter size across domestic dog breeds, we conducted an across-breed GWAS analysis using 115,683 SNPs and litter size data from 60 dog breeds (Figure IB, Supplementary Figure 2B). We identified two significant and one marginally significant SNPs (Supplementary Table S4) intersecting three genes, namely keratin 71 (*KRT71*, uncorrected *P* = 2.2 x 10^−8^, *Perm P* = 0.017), RNA Terminal Phosphate Cyclase-Like 1 *(RCL1*, uncorrected *P* = 2.6 x 10^−8^, *Perm P—* 0.001) and microphthalmia-associated transcription factor (*MITF*, uncorrected *P* = 3.5 x 10^−7^, *Perm P* = 0.079). The *KRT71* SNP is the same variant that was marginally associated with variation in cesarean section rate described above, but in this instance its association with the trait remains significant after permutation analysis. Another three significant SNPs were found in intergenic regions; two were nearby genes *MSRB3* (methionine sulfoxide reductase B3, uncorrected *P* = 1.3 x 10^∼7^, *Perm P—* 0.001) and *MSANTD1* (Myb/SANT DNA binding domain containing, uncorrected *P* = 1.5 x 10^−9^, *Perm P* = 0.001), respectively. The final variant was near an RNA of unknown function (ENSCAFG00000021196, uncorrected *P* = 3.8 x 10^−10^, *Perm P* = 0.001).

The *RCL1* SNP (chromosome 1: 93,189,363) is found in the intron between exons 7 and 8. *RCL1* functions in the maturation of 18s RNA (Lyng et al. 2006) and is associated with cervical cancer; one role of the gene in this cancer pathology is thought to involve the regulation of insulin receptors (Lyng et al. 2006). Additionally, a rare missense variation in *RCL1* was recently associated with depression (Amin et al. 2017).

The *KRT71* SNP (chromosome 27: 2,539,211) results in a missense mutation of exon 2 of the gene, which belongs to a family of keratin genes specifically expressed in the inner root sheath of hair follicles (Langbein et al. 2003). Prior analysis in dogs identified variation in gene *KRT71*, along with variation in genes *RSP02* and *FGF5*, accounting for most coat phenotypes (Cadieu et al. 2009), such as curliness.

Another SNP (chromosome 10: 8,114,328) significantly associated with litter size is found in the intergenic region downstream of *MSRB3*, whose protein product catalyzes the reduction of methionine-R-sulfoxides to methionine and repairs oxidatively damaged proteins (Brot et al. 1981, Kim et al. 2004). In humans, mutations in *MSRB3* are associated with deafness (Ahmed et al. 2011). Epigenetic changes of *MSRB3* in the fetus during pregnancy may affect length of gestation, with increased DNA methylation correlated with increased gestational age (Lee et al. 2012, Schierding et al. 2014). Furthermore, *MSRB3* shows an increase in mRNA expression in ripe (at term) versus unripe human uterine cervix, implying that *MSRB3* functions to ripen the cervix before the onset of labor (Hassan et al. 2009). In previous morphological studies in dogs, *MSRB3* is associated with ear erectness (Boyko et al. 2010).

The last SNP (chromosome 6: 61,062,626) that is significantly associated with litter size is located downstream of *MSANTD1*, which is part of a gene network believed to aid in cell-to-cell signaling and interaction, hematological system development and function, and immune cell trafficking (Yu et al. 2014). *MSANTD1* has been identified in two independent studies as a candidate gene for the determination of black coat color in goats (Benjelloun et al. 2015, Wang et al. 2016).

Finally, following permutation analysis, the association between the *MITF* SNP (*Perm P =* 0.079) and litter size was just above the threshold for significance and can only be considered as suggestive. The *MITF* SNP (chromosome 20: 21,848,176) is found in the intron between exons 4 and 5. *MITF* plays an integral role in the development of neural crest-derived melanocytes and optic cup-derived retinal pigment epithelial cells. In human melanocytes, *MITF* is a regulator of *D1APH1*, a member of the formin gene family whose members are highly expressed in reproductive tissues and have been associated with a variety of reproductive phenotypes (Carreira et al. 2006, Lamm et al. 2018, Cruickshank et al. 2013, Elovitz et al. 2014, Montenegro et al. 2009). *D1APH1* expression is increased in spontaneous term and preterm labor myometrial tissues (Lartey et al. 2007). In domesticated animals, *MITF* is a well characterized gene associated with coat color (Boyko et al. 2010, Wang et al. 2016). In humans, *MITF* is expressed in melanocytes, as well as reproductive tissues including the endometrium and cervix (Figure 2) (Uhlén et al. 2015).

### Genetic loci that significantly associate with stillbirth rate

To examine whether there are SNPs that are significantly associated with variation in stillbirth rate among breeds, we retrieved data for stillbirth rates for 57 of the 162 dog breeds (Supplementary Table 1). The data covers 10,810 litters of 224 breeds registered in the Norwegian Kennel Club and defines perinatal mortality as the sum of stillborn puppies and puppies that died during the first week after birth (Tønnessen et al. 2012). Among these 57 breeds with overlapping genomic data, the median stillbirth rate is 4.2 pups, with a maximum rate of 12.3% in Saint Bernards and a minimum of 0% in Basenjis and Italian Greyhounds (Supplementary Figure 3C).

To test if any SNPs are significantly associated with the observed variation in stillbirth rate across domestic dog breeds, we conducted an across-breed GWAS analysis using 115,683 SNPs and stillbirth rate data from 56 dog breeds (Figure 1C, Supplementary Figure 2C). We identified five significant and marginally significant SNPs (Supplementary Table S5); five intersecting 4 genes, 5-Hydroxtryptamine receptor 2C (*HTR2C*, uncorrected *P* = 2.0 x 10^−7^, *Perm P* = 0.001), keratin 71 *{KRT71*, uncorrected *P* = 3.2 x 10^−9^,*Perm P* = 0.064), and microphthalmia-associated transcription factor (*MITF*, uncorrected *P* = 1.4 x 10^−7^, *Perm P* = 0.079), and SP140 nuclear body protein (*SP140*, uncorrected *P* = 2.76 x 10^−8^, *Perm P* = 0.001) and one in an intergenic region near a snoRNA (ENSCAFG00000027305, uncorrected *P* = 1.3 x 10“^7^, *Perm P =* 0.002) of unknown function. The *KRT71* SNP associated with variation in stillbirth rate is the same one as that associated with litter size described above. Similarly, the *MITF* SNP associated with variation in stillbirth rate is the same as that associated with litter size. However, both of these associations have permutation p-values > 0.05 and thus can only be considered as tentative.

The *SP140* SNP (chromosome 25: 42,482,266) resides in the intro between exons 4 and 5. *SP140* is the lymphoid-restricted homolog of *SP100* expressed in mature B cells, as well as some T cells (Bloch et al. 1996). High levels of *SP140* mRNA are detected in human spleen and peripheral blood leukocytes, but not other human tissues (Bloch et al., 1996). *SP140* expression has been implicated in innate response to immunodeficiency virus type 1 (Madani et al. 2002). Finally, *SP140* was the gene showing the largest difference in expression level between normal and preeclamptic placentas (Heikkilä et al. 2005).

The *HTR2C* SNP (chromosome X: 87,378,551) is located in the intron between exons 3 and 4. *HTR2C* is one of the most important and extensively studied serotonin receptors (Drago et al. 2009). *HTR2C* has ten fixed SNP differences between dogs and wolves, and also belongs to the behavioral fear response (Li et al. 2013). Additionally, *HTR2C* is differentially expressed in the brain between tame and aggressive mice and foxes (Kukekova et al. 2011), providing additional evidence for its involvement in the tame behaviors of domesticated dogs (Li et al. 2013).

### Genetic loci that significantly associate with gestation length

To examine whether there is variation in gestation length among breeds, we identified individual gestation length averages by breed predominantly in breeder handbooks. Utilizing breeders’ handbooks, we were able to identify gestation length means for a total of 23 of the 162 dog breeds that we had genotypic data for (Supplementary Table 1). Among these 23 breeds, the median gestation length is 62.2 days, with a maximum length of 65.3 in beagles and a minimum of 60.1 in the Alaskan Malamute (Supplementary Figure 3D).

To identify SNPs, and genes proximal to them, that are significantly associated with the observed variation in gestation length across domestic dog breeds, we conducted an across-breed GWAS analysis using 115,683 SNPs and gestation length data from 23 dog breeds (Figure ID, Supplementary Figure 2D). Our analysis identified six significantly associated SNPs (Supplementary Table S6) that mapped to 4 genes, namely solute carrier family 9 (*SLC9A8*, uncorrected *P* = 3.7 x 10^−11^, *Perm P* = 0.001), calcium channel, voltage-dependent, alpha-2/delta Subunit 3 (*CACNA2D3*, uncorrected *P* = 3.1 x 10^−7^, *Perm P* = 0.013), microtubule associated tumor suppressor candidate 2 (*MTUS2*, uncorrected *P* = 3.6 x 10^−7^, *Perm P* = 0.001), and helicase family member 1 (*HFM1*, uncorrected *P* = 4.0 x 10^−7^, *Perm P* = 0.013), and two lincRNAs (ENSCAFG00000037743, uncorrected *P* = 4.4 x 10^−7^, Perm *P* = 0.001, and ENSCAFG00000039067, uncorrected *P* = 1.6 x 10^−7^, Perm P = 0.001) whose functions are unknown.

The first significantly associated SNP (chromosome 24: 36,399,705) resides in intron 78 of *SLC9A8*, an integral transmembrane protein that exchanges extracellular Na+ for intracellular H+. *SLC9A8* serves multiple functions, including intracellular pH homeostasis, cell volume regulation, and electroneutral NaCl absorption in epithelia (Xu et al. 2008). Knockout male mice have impaired luteinizing hormone-stimulated cAMP production and are infertile, despite normal morphology of their reproductive system and normal behavior (Xu et al. 2015). *SLC9A8* is expressed ubiquitously (Figure 2) (Uhlén et al. 2015), an expression pattern suggestive of involvement in housekeeping functions.

The second SNP (chromosome 20: 35,206,774) is found in the intron between exons 26 and 27 of *CACNA2D3*. This gene is one of four members of the alpha-2/delta subunit three family of the voltage-dependent calcium (Ca2+) channel complex, regulating the influx of Ca2+ ions entering the cell upon membrane polarization (Jin et al. 2017). The regulation of calcium is a fundamental process relevant to life at fertilization, and subsequent control of development and differentiation of cells (Berridge et al. 1998). In previous studies in humans, *CACNA2D3* is differentially methylated in the amnion between normal and preeclamptic pregnancies (Suzuki et al. 2016) and in blood between extreme preterm and term infants at birth (Cruickshank et al. 2013, Eidem et al. 2015). Additionally, *CACNA2D3* is one of four genes recently described as influencing cranial morphology in human populations (Paternoster et al. 2012). In other domesticated animals, *CACNA2D3* is downregulated by Colony Stimulating Factor 2 (*CSF2*) in the trophoectoderm of pregnant cattle, which increases the ability of the preimplantation embryo to advance to the blastocyst stage (Ozawa et al. 2016). In the closely related wolf, *CACNA2D3* is under diversifying selection associated with environmental adaptations to altitude (Pilot et al. 2014, Schweizer et al. 2016, Zhang et al. 2014).

The third significantly associated SNP (chromosome 25:10,481,606) falls in a large intronic region of the *MTUS2* gene. The protein product of *MTUS2* is cardiac zipper protein (CAZIP), a member of a class of proteins that interact with angiotensin II receptor interacting proteins (ATIP) (Rodrigues-Ferreira et al. 2010). *MTUS2* plays a role in the development and function of the heart and nervous system in vertebrates (Puy et al. 2009).

The fourth SNP (chromosome 6: 57,457,184) is located in the 3’ intron of *HFM1*, a DNA helicase that confers genome integrity in germline tissues (Tanaka et al. 2009). *HMF1* plays a role in meiotic recombination implying a major evolutionary role through the creation of diverse offspring. In mice, deletion of *HFM1* eliminates a major fraction of cross over events (Guiraldelli et al. 2013), whereas in cattle *HMF1* is associated with both fertility and milk production in Holstein cattle (Pimentel et al. 2011), as well as with alteration of global recombination rates in Holstein, Holstein-Friesian, Jersey, and crossbred individuals (Kadri et al. 2016).

## Discussion

Mammals exhibit a great deal of variation in their reproductive traits, yet remarkably little is known about the genetic basis of these traits. To begin to address this, we used GWAS analyses to examine the genetic basis of four reproductive traits (cesarean section rate, stillbirth rate, litter size, and gestation length) across up to 97 domestic dog breeds. We identified several significant genetic associations for each trait (Figure 1).

Five of the 13 genetic variants that we found to be associated with reproductive trait variation have been previously identified to be involved in diverse traits associated with dog domestication (Table 2), such as brachycephaly and coat curl and color, suggesting that selection for signature traits of dog breeds may have also directly or indirectly influenced variation in reproductive traits. For example, one of the variants that we found to be associated with cesarean section rate is in an intron of *SM0C2*, a gene previously associated with brachycephaly in dogs (Bannasch et al. 2010, Marchant et al. 2017). Brachycephaly, the shortening and widening of the muzzle and skull, is present in several “fighting” breeds such as Boxer, Boston Terrier, and Bulldog, and is thought to have been originally artificially selected on the basis that a shorter and wider cranial shape would enhance the dog’s biting power (Ellis et al. 2009). Interestingly, one of the traits that associated with brachycephaly is cephalopelvic disproportion (Evans and Adams 2010), a significant medical condition that can result in the death of both the litter and the bitch due to the inability of the pups to pass through the pelvic canal. The negative effects of cephalopelvic disproportion are alleviated by cesarean section, which not only allows these breeds to reproduce but also enables the continued application of artificial selection for the most extreme cranial morphology (Bannasch et al. 2010). Whether the *SMOC2* variant identified directly influences parturition and birth timing in dogs (in humans, *SMOC2* is highly expressed in several reproductive tissues; see Figure 2 and Ref. (Uhlén et al. 2015) or indirectly leads to adverse pregnancy outcomes (e.g. brachycephalic cranial morphology leading to cesarean section) remains unknown. It is highly likely, however, that the association between *SMOC2* and brachycephaly came first, paving the way for the subsequent association of both with cesarean section rate.

Several of the significantly associated genes that we identified in dogs appear to also be associated with reproductive phenotypes in humans. This suggests the possibility that the artificial selection that gave rise to dog breeds may have also contributed to the observed variation in their reproductive traits. For example, a member of the gene family for a subunit of the voltage-dependent calcium channel complex, *CACNA2D3*, which is associated with gestation length in our study, has been shown to be both differentially methylated in amnion between normal and preeclamptic human pregnancies (Suzuki et al. 2016), and in blood between extreme preterm and term infants at birth (Cruickshank et al. 2013, Eidem et al. 2015). Furthermore, expression of *MSRB3*, which is associated with litter size in our study, is elevated in ripe (at term) versus unripe human uterine cervix and may be involved in the onset of labor (Hassan et al. 2009). Finally, a few of the other genes significantly or marginally associated with reproductive traits (*SMOC2* and *MITF)* are also known to be expressed in human reproductive tissues (Uhlén et al. 2015)(Figure 2).

## Methods

### Genotypic and Phenotypic Data

To identify SNPs that are significantly associated with reproductive traits, we used a previously published data set containing 160,727 SNPs from 4,342 individual dogs across 162 breeds genotyped using the Illumina 173k CanineHD array that were downloaded from http://datadryad.Org/resource/doi:10.5061/dryad.266k4 (Hayward etal. 2016). Following the original authors, SNPs with a genotyping rate (i.e., the proportion of genotypes per marker with non-missing data) below 95% and heterozygosity ratios (i.e., the ratio of the number of heterozygous SNPs divided by the number of non-reference homozygous SNPs) below 0.25 or above 1.0 were removed.

Phenotypic reproductive trait data for litter size (number of pups), cesarean rate, stillbirth rate, and gestation length across 128 breeds were collected from a variety of breeder’s handbook and primary journal articles (Borge et al. 2011, Tonnessen etal. 2012, Evans and Adams 2010, Chatdarong et al. 2007, Concannon et al. 1983, Elits et al. 2005, Evans and White 1997, Gavrilovic et al. 2008, Okkens et al. 1993, Son et al. 2001, Kim et al. 2007, Linde-Forsberg et al.) (see also Supplementary File 1). We also included body mass as a control trait. Each breed was assigned the average breed value for each phenotype; the full list of the values for all four reproductive traits and body mass across the 128 breeds is provided in supplementary table 1. For the body mass control, our collected trait values overlapped with the genotypic data (Hayward et al. 2016) for 101 breeds corresponding to 3,384 individuals (Table 1). For the reproductive traits, our collected cesarean section rate trait values overlapped with the genotypic data for 95 breeds (3,194 individuals), our litter size trait values for 60 breeds (2,617 individuals), our stillbirth rate values for 56 breeds (2,590 individuals), and our gestation length values for 23 breeds (1,908 individuals) (Table 1).

**Table 1.**
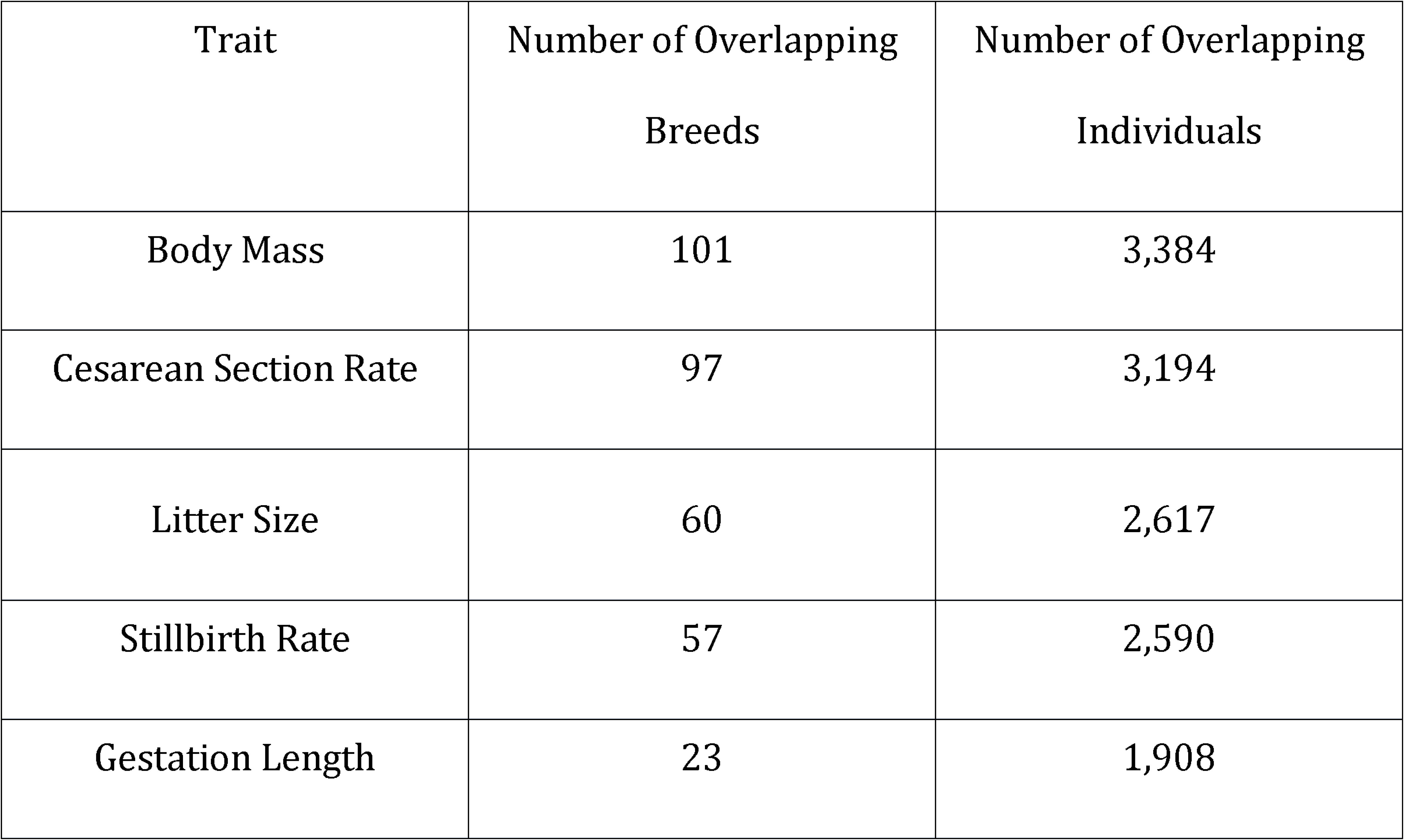
Numbers of breeds and individuals with overlapping phenotypes and genotypes included in our analysis.

| Trait | Number of Overlapping Breeds | Number of Overlapping Individuals |
| --- | --- | --- |
| Body Mass | 101 | 3,384 |
| Cesarean Section Rate | 97 | 3,194 |
| Litter Size | 60 | 2,617 |
| Stillbirth Rate | 57 | 2,590 |
| Gestation Length | 23 | 1,908 |

### Genome Wide Association (GWAS) Analyses

To test SNPs for associations with the four reproductive traits of interest, we conducted a GWAS analysis for each individual trait using body mass as a covariate, and accounting for kinship, as well as for body mass as a proof of concept. All GWAS analyses were run using a linear-mixed model as implemented in the program GEMMA, version 0.94 (Zhou and Stephens 2012). Numerous studies have shown that the vast majority of morphological, ecological and physiological traits vary as a function of an organism’s body mass (Gould 1966, Kleiber 1932, Shingleton 2010) as well as a function of kinship (Hayward et al. 2016, Boyko et al. 2010). Most notably for the purpose of this study, body mass has been previously shown to be strongly correlated with litter weight (Sacher and Staffeldt 1974, Blueweiss et al. 1978, Tuomi 1980), neonate weight (Sacher and Staffeldt 1974, Blueweiss et al. 1978, Tuomi 1980, Ross 1988), and gestation length (Martin et al. 1985, Sacher and Staffeldt 1974, Blueweiss et al. 1978, Kihlström 1972, Phillips etal. 2015).

To ensure our analysis reflected the reproductive trait of interest and not SNPs associated with body mass, we used log body mass as a covariate for all reproductive trait analyses. To be able to do so, we pruned our genotypic data so that they included only dog breeds (and individuals) for which we had both body mass and reproductive trait of interest values (see Supplementary Table 1).

To account for population stratification, we calculated a kinship matrix of the included breeds using GEMMA and included it as a random effect in each association analysis. Each value of a kinship matrix describes the probability that a particular allele from two randomly chosen individuals at a given locus is identical by descent (Lange 2002). Finally, to control for inflated P value significance from the testing of multiple hypotheses, we used a significance threshold of *P* = 4.3 x 10^_7^(Bonferroni cutoff of *α* = 0.05, *N* = 115,574) for all analyses. All reported P values are Wald’s P values as calculated in GEMMA (Zhou and Stephens 2012).

To reduce potential error stemming from SNP misidentification in our analyses, we included only SNPs with a minor allele frequency (MAF) > 0.05, since SNPs with very low minor allele frequencies are more prone to error due to the small number of samples that have the called nucleotide. Furthermore, we pruned SNPs not in complete or near-complete linkage disequilibrium using a variance inflation factor of 10, using the PLINK command - indep 100 10 10 (Purcell etal. 2007).

Finally, to validate variants significantly associated with at least one reproductive life history trait in the domestic dog, we performed a permutation analysis. For each trait, we randomly permuted the assignment of breed-specific phenotypes while holding body mass constant for each breed across 1,000 permutations. We then regressed the randomly assigned reproductive phenotypes onto log body mass and assigned the residual for each breed as the phenotype for all of the individuals of that breed. Next, using GEMMA (Zhou and Stephens 2012), we performed an association test for each variant to obtain a single permuted p-value. The permutation p-value in Table 2 corresponds to the number of times that a single permuted p-value was less than the empirical p-value from the original analysis. Any variant with a permutation p-value greater than 0.05 was acknowledged as a potential false positive association.

**Table 2.**
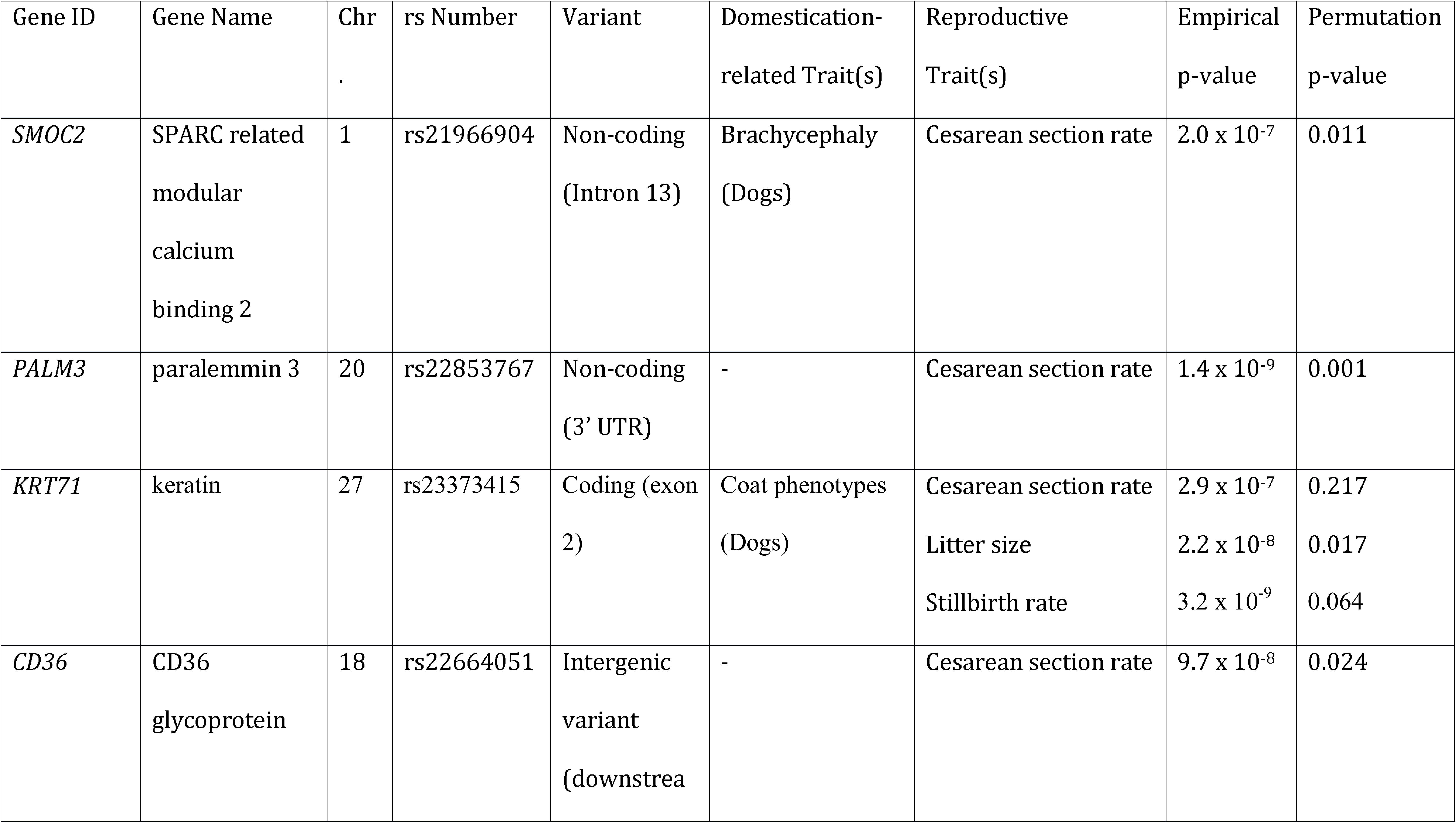
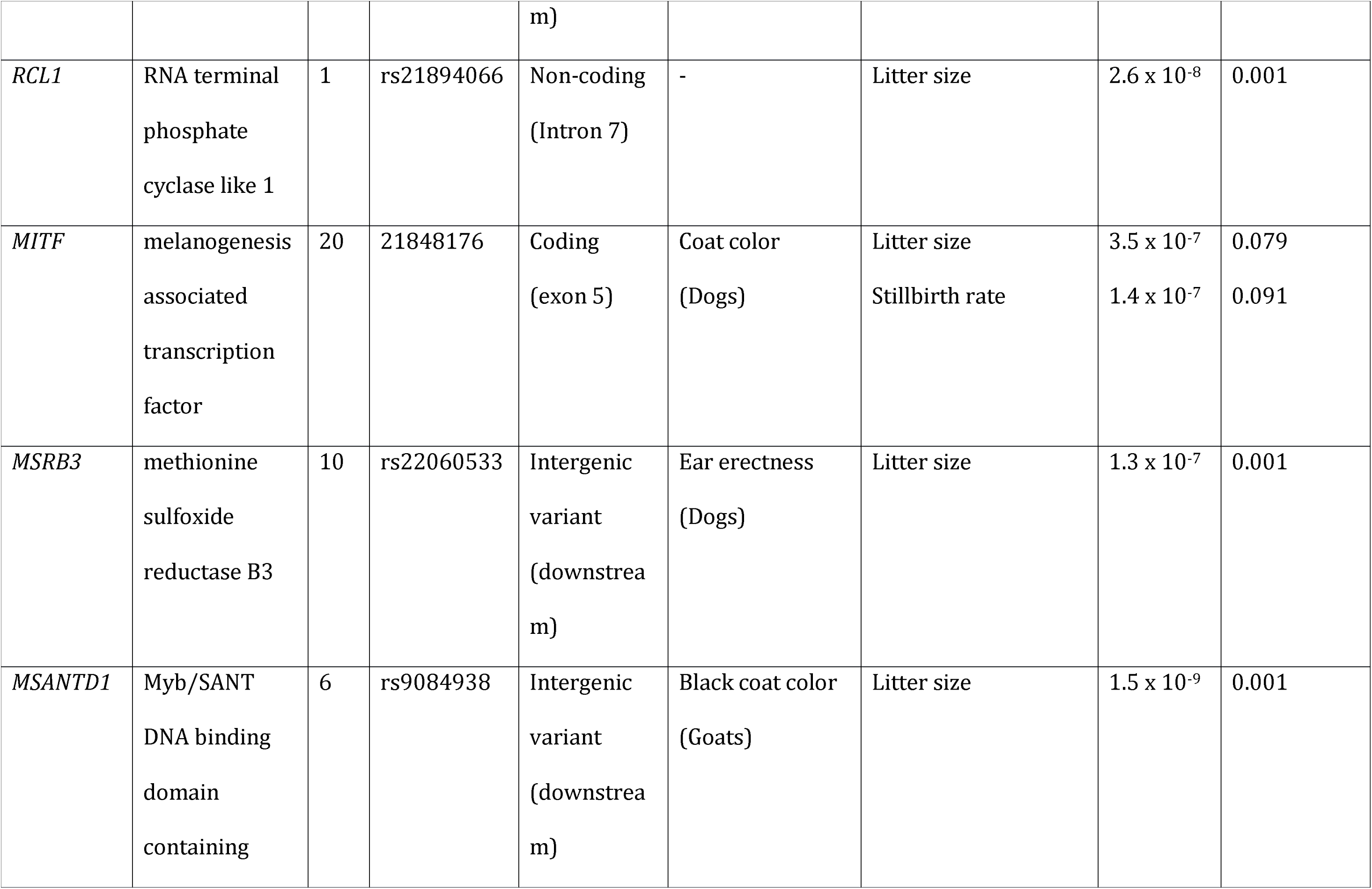
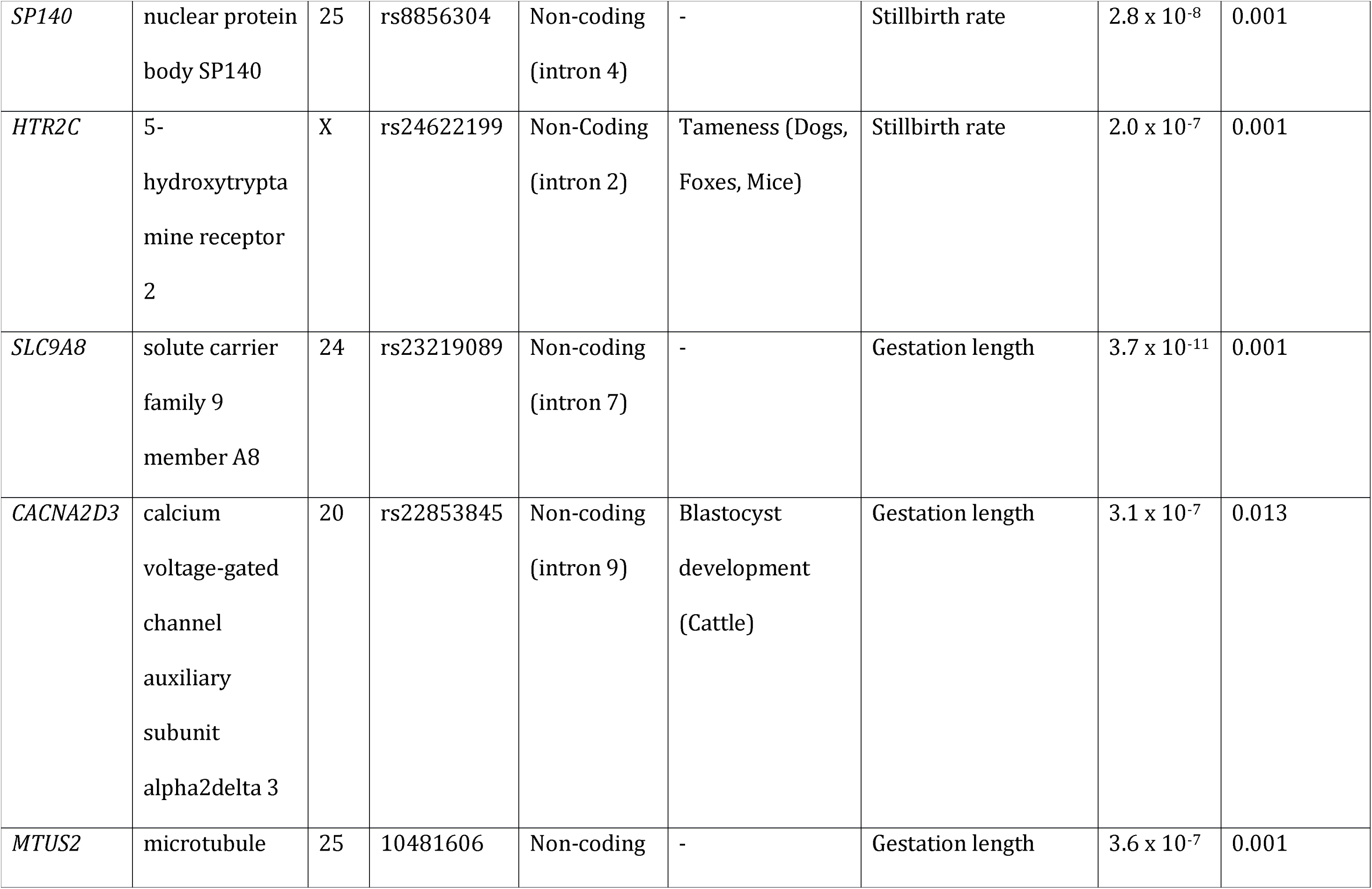
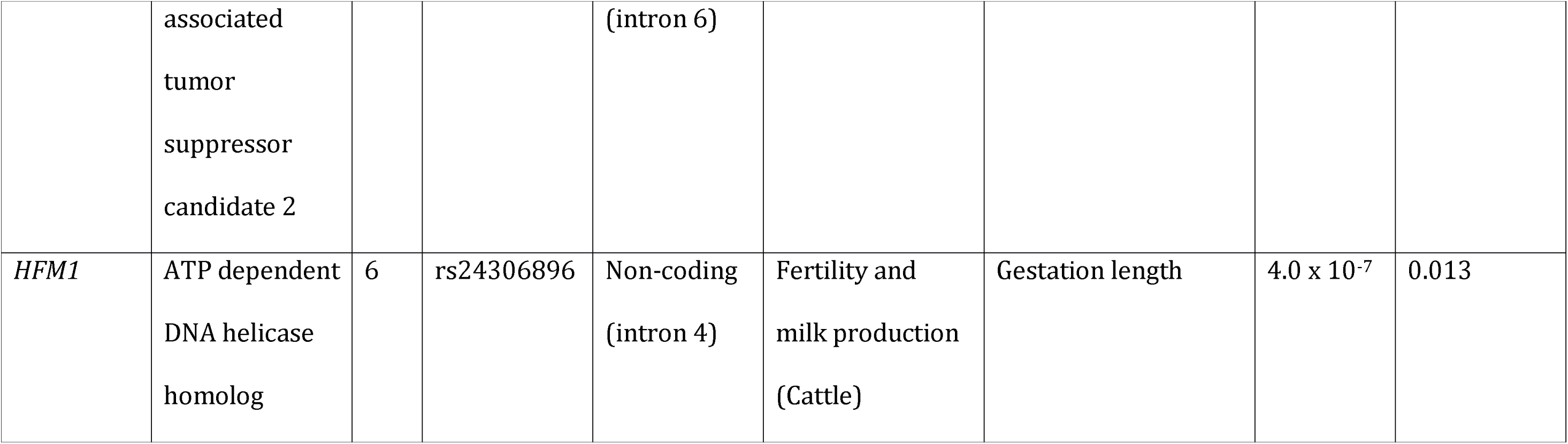
Summary of genes that contain or are adjacent to the SNPs that are associated with variation in reproductive traits across dog breeds.

To gain insight into the genetic elements putatively involved with the traits of interest, we mapped all SNPs found to be significantly and marginally associated with each trait of interest using custom perl and R scripts to the CanFam3.1.87 dog genome assembly (Lindblad-Toh et al. 2005, Hoeppner et al. 2014). Transcript IDs were mapped to gene names using bioconductor biomaRt interface to the ENSEML biomart (Durinck et al. 2009). If the significant SNP was outside gene boundaries, we reported the nearest upstream or downstream gene. Manhattan plots and quantile-quantile plots were generated using R 3.1.2 (R Core Team 2013) with the qqman package (Turner 2014). Calculation of the *X* inflation parameter, a metric of any existing systematic bias in the data set, was calculated using the GenABEL R package (Aulchenko 2007) and was used to interpret Type I error rate in the multiple testing of GWAS analyses (Rao et al. 2016).

## Acknowledgements

We thank members of the Rokas lab and members of the March of Dimes Prematurity Research Center Ohio Collaborative for useful discussions of this work.

## Funding Statement

This research was supported by the March of Dimes through the March of Dimes Prematurity Research Center Ohio Collaborative (to A.R., P.A. and J.A.C) and by the Burroughs Wellcome Fund (to J.A.C and A.R.). MLJ received partial support through a Littlejohn Summer Research Scholarship at Vanderbilt University. The funders had no role in study design, data collection and analysis, decision to publish, or preparation of the manuscript.

## Supplementary Material

**Supplementary Figure 1. Recapitulation of SNPs associated with body mass in 101 domesticated dog breeds.** (A) Body mass distribution for 101 breeds. (B) Manhattan plots showing the statistical significance of each SNP as a function of genomic position for body mass. Plot generated in R using the qqman package. (C) Quantile-quantile plot showing the effectiveness of the stratification correction (λ = 1.17). Plot generated in R; inflation factor was calculated using the GenABEL package implemented in R.

**Supplementary Figure 2. Distribution of phenotypic values of the four reproductive traits examined in this study across dog breeds.** (A) cesarean section rate (n = 97 breeds), (B) litter size (n = 60 breeds), (C) stillbirth rate (n = 57 breeds), and (D) gestation length (n = 23 breeds). Plots were generated in R using the gglplot2 package.

**Supplementary Figure 3. Quantile-quantile plots for the GWAS analyses of the four reproductive traits.** The range for the inflation factor (λ) for all GWAS analyses is between 1.05 – 1.09, indicating the effectiveness of the stratification correction. (A) cesarean section rate (λ = 1.05), (B) litter size (λ = 1.05), (C) stillbirth rate (λ = 1.05), and (D) gestation length (λ = 1.09). Plots generated in R, and inflation factors were calculated using the GenABEL package implanted in R.

**Supplementary Table 1. Summary of raw phenotypes for breeds included in analysis.**

**Supplementary Table 3. Summary of top 100 SNPs associated with cesarean section rate.**

**Supplementary Table 4. Summary of top 100 SNPs associated with litter size. Supplementary**

**Table 5. Summary of top 100 SNPs associated with stillbirth rate.**

**Supplementary Table 6. Summary of top 100 SNPs associated with gestation length.**

**Supplementary File 1. Sources of phenotypic data.**

